# Caveats and nuances of model-based and model-free representational connectivity analysis

**DOI:** 10.1101/2021.08.10.455841

**Authors:** Hamid Karimi-Rouzbahani, Alexandra Woolgar, Richard Henson, Hamed Nili

**Affiliations:** MRC Cognition & Brain Sciences Unit, University of Cambridge, UK; Department of Computing, Macquarie University, Australia; Department of Psychiatry, University of Cambridge, UK; Wellcome Centre for Integrative Neuroimaging, University of Oxford, UK

**Keywords:** representational connectivity analysis, multi-dimensional connectivity, functional connectivity, multivariate pattern analysis, representational similarity analysis

## Abstract

Brain connectivity analyses have conventionally relied on statistical relationship between one-dimensional summaries of activation in different brain areas. However, summarising activation patterns within each area to a single dimension ignores the potential statistical dependencies between their multi-dimensional activity patterns. Representational Connectivity Analyses (RCA) is a method that quantifies the relationship between multi-dimensional patterns of activity without reducing the dimensionality of the data. We consider two variants of RCA. In model-free RCA, the goal is to quantify the shared information for two brain regions. In model-based RCA, one tests whether two regions have shared information about a specific aspect of the stimuli/task, as defined by a model. However, this is a new approach and the potential caveats of model-free and model-based RCA are still understudied. We first explain how model-based RCA detects connectivity through the lens of models, and then present three scenarios where model-based and model-free RCA give discrepant results. These conflicting results complicate the interpretation of functional connectivity. We highlight the challenges in three scenarios: complex intermediate models, common patterns across regions and transformation of representational structure across brain regions. The paper is accompanied by scripts that reproduce the results. In each case, we suggest potential ways to mitigate the difficulties caused by inconsistent results. The results of this study shed light on some understudied aspects of RCA, and allow researchers to use the method more effectively.

## 1. Introduction

To study the neural underpinnings of cognitive processes, we need not only to characterise the response of individual brain regions, but understand the functional connectivity between them. This is critical to understand how brain regions interact in giving rise to cognition (Bressler and Menon, 2010). Functional connectivity across the brain has been conventionally evaluated using univariate/one-dimensional analyses (Bastos and Schoffelen, 2016). In these analyses, responses in different brain regions are initially summarized for each region in a one-dimensional metric (Biswal et al., 1995). If these metrics are statistically-related, we then infer functional connectivity between the regions (Bastos and Schoffelen, 2016). For example, methods such as gamma-band synchronization (Gregoriou et al., 2009), phase covariance across regions (Bar et al., 2006), frequency coupling (Karimi-Rouzbahani et al., 2021b) and differential equations (Friston et al., 2013) have been used to evaluate connectivity after summarizing the activation patterns across vertices (sensors or voxels) within each region. However, univariate connectivity analysis can miss connectivity if the pairs of regions are statistically related through multidimensional patterns of activation rather than the summarized (e.g. averaged) activation within each region (Coutanche, 2013; Basti et al., 2019; Basti et al., 2020). For example, for heterogeneous ROIs, where multiple response modes co-exist, projecting multivariate response patterns onto one dimension could lead to strong distortions. This has led to a recent shift from univariate to multidimensional (multivariate) connectivity analyses (Coutanche and Thompson-Schill, 2013; Goddard et al., 2016; Anzellotti and Coutanche, 2018; Basti et al., 2019; Basti et al., 2020; Karimi-Rouzbahani et al., 2021a; Karimi-Rouzbahani et al., 2021c; Shahbazi et al., 2021). One approach to multidimensional connectivity analyses is Representational Connectivity Analysis (RCA; Kriegeskorte et al., 2008), which utilizes the versatility of Representational Similarity Analysis (RSA) to move from the direct comparison of representations to the comparison of representational geometries (Kriegeskorte et al., 2008). RCA can be divided into model-free and model-based methods, each having specific characteristics. Here, we describe model-free and model-based RCA and point out their differences. Specifically, we present three simple scenarios where model-free and model-based RCA provide inconsistent connectivity results, flagging the situations where they should be used with caution and adding nuance to how the results of each should be interpreted.

One key feature of RCA is that, rather than activations (Anzellotti et al., 2017a; Anzellotti et al., 2017b; Basti et al., 2019), it evaluates the statistical dependency between the geometry/structure of neural representations across areas. Accordingly, RCA relies on the distinctiveness (i.e. dissimilarity) of patterns across conditions, which is conceived in terms of “information encoding/representation”, rather than the activity patterns themselves. Therefore, one prerequisite for performing RCA is to have enough distinct experimental conditions to obtain the geometry of representations in the neural data (see, however, how we performed RCA on a single condition across the trial (Karimi-Rouzbahani et al., 2021c)). This usually precludes RCA from being used to test functional connectivity in resting-state data (single, continuous fMRI or M/EEG time series), which dominates univariate functional connectivity analyses. On the flip side, however, the representational nature of RCA provides several advantages over activity-based connectivity analyses. First, RCA allows the evaluation of connectivity across any two regions with different number of response channels (i.e. vertices, voxels, sensors or sources; Kriegeskorte et al., 2008). Second, it allows one to ask how information (e.g. sensory, cognitive, etc.), rather than activation, is potentially transferred across areas. Third, model-based RCA allows one to target specific aspects of information, based on hypotheses about how a specific aspect of information is transferred, and avoiding the influence from undesired confounders on connectivity (Clarke et al., 2018; Karimi-Rouzbahani et al., 2021a). Despite these advantages, under some circumstances representational connectivity analysis can miss true connectivity or erroneously detect non-existing (false) connectivity. This necessitates further investigation of RCA methods before they are more widely used as measures of multi-dimensional connectivity.

From a broad perspective, model-free RCA (Basti et al., 2020; Coutanche et al., 2020; Karimi-Rouzbahani et al., 2021c; Shahbazi et al., 2021) evaluates whether there is any commonality in the distributed patterns of activity for two brain regions. The commonality might reflect shared information due to similar encoding in the two regions. But it might also be due to the encoding of nuisance factors that are shared across regions. On the contrary, model-based RCA asks whether the two regions have shared information with regards to a specific hypothesis defined by a model. Having the model(s) can give a more specific picture of the multi-dimensional regional interactions. In this paper, we compare model-free and model-based RCA and explain their pros and cons. In particular, we raise some cautions for using each method by showing simulated cases where one method fails to capture functional connectivity between two regions with shared information.

## 2. Methods and Results

### Simulations

we generated multidimensional patterns of activity using scripts from the Matlab RSA toolbox (Nili et al., 2014). The Matlab script for reproducing the results can be downloaded from https://osf.io/3nxfa/. We simulated activity patterns for 16 stimuli in two brain regions. The 16 conditions can be thought of as corresponding to 4 peripheral positions of the visual field (e.g. top left, top right, bottom left and bottom right) of 4 semantically distinct visually presented object categories (e.g. animals, faces, fruits and objects). For simpler explanation and interpretation of the results one can think of region of interest (ROI) 1 as visual area 2 (V2) and ROI 2 as inferior temporal cortex (ITC). Accordingly, ROI 1 dominantly represents position (i.e. regardless of the category of the objects) and ROI 2 dominantly represents semantic categories (i.e. regardless of the position of the stimuli). Figure 1A depicts the arrangements of the conditions in the Representational Dissimilarity Matrix (RDM), and Figure 1B shows the ground-truth of the RDMs in the two ROIs and neural RDMs for a simulated subject which are noisier versions of the ground truth.

**Figure 1.**
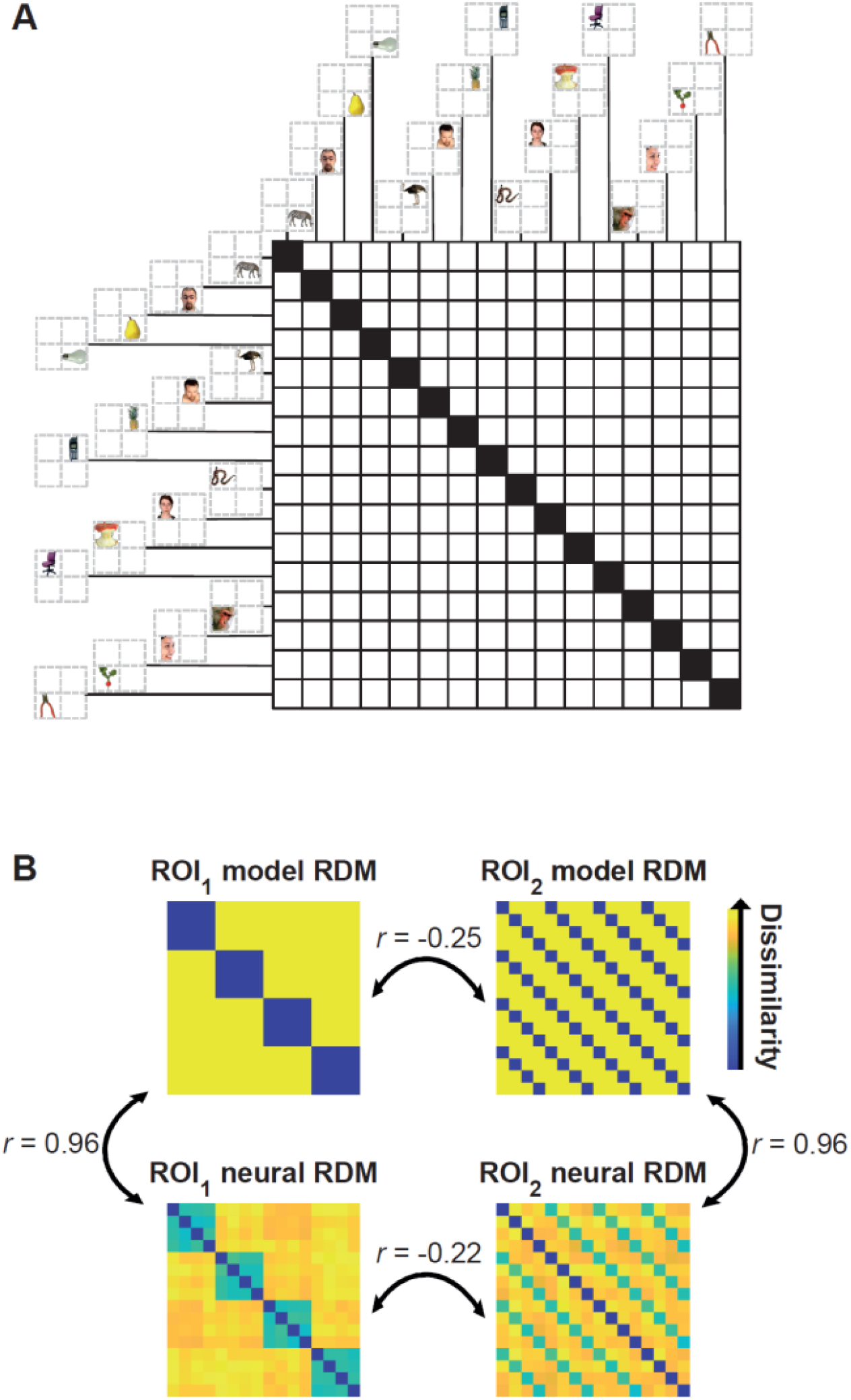
Arrangement of the simulated conditions in the representational dissimilarity matrices (RDMs), model and neural RDMs and their correlations. (A) There are 16 conditions in the RDM consisting of four semantic categories of objects in four distinct positions. Stimuli are from Kriegeskorte et al., 2008 available at http://www.cns.nyu.edu/kianilab/Datasets.html. (B) Top matrices show the ground truth for the ROI that represents positions (left) and semantic categories (right). The neural RDMs shown in the bottom were generated by adding noise to the activity patterns used in the two top model RDMs. Therefore, the simulated neural RDMs are highly correlated to the ground truth (r > 0.95). The correlations between the (model and/or neural) RDMs of the two different ROIs are negative (r < −0.2) implying no connectivity between them.

We used Pearson’s (linear) correlation for comparing RDMs. Accordingly, we only considered significantly positive correlations as indicating representational connectivity. We performed significance testing using a one-sided Wilcoxon’s signed rank test (Wilcoxon, 1992) across subjects and applied a threshold of 0.001 for statistical significance. Note that as RDMs are symmetric matrices, we only analysed the elements in the upper triangle excluding the diagonal. We simulated data for N = 20 subjects to match it to the conventional number of participants in real-life neuroimaging experiments and performed the statistical tests at group level.

### 2.1 Simulation 1: model-based RCA tests connectivity through the lens of model(s)

Having hypotheses when testing for representational connectivity allows us to test whether two ROIs are related with regards to a specific model^1^. A model privileges a specific direction in the dissimilarity space so that all comparisons would be made with respect to the direction specified by the model. One can think of different ways to implement model-based RCA (Clarke et al., 2018; Karimi-Rouzbahani et al., 2021a). Here, we rely on the concept that model-based RCA evaluates the extent to which a model contributes to the representations of two ROIs. Therefore, we use a minimalistic implementation of model-based RCA to deliver some caveats as clearly as we can.

Despite the benefits offered by model-based RCA, it has some properties that should be considered with caution. For example, consider the scenarios depicted in Figure 2A. Although RDMs reside in a high dimensional dissimilarity space, we illustrate the main point in 2D figures. Figure 2A shows a case where two ROIs have a positive correlation to a model RDM and in fact identical similarities to it (e.g. Pearson correlation of 0.7). Conceptually, model-based RCA asks whether correlations of two brain RDMs to a model RDM are close or not, so the two ROIs are considered connected according to the model. However, the RDMs in the two ROIs are orthogonal. Therefore, unlike model-based RCA, model-free RCA would conclude no functional connectivity. Conversely, consider the case depicted in Figure 2B. The neural RDMs in the two ROIs have a positive correlation (e.g. a Pearson correlation of 0.7). This means that model-free RCA would indicate representational connectivity. However, one ROI has no relationship to the model (correlation of 0), and therefore from the perspective of the model, the two ROIs would not be connected.

**Figure 2.**
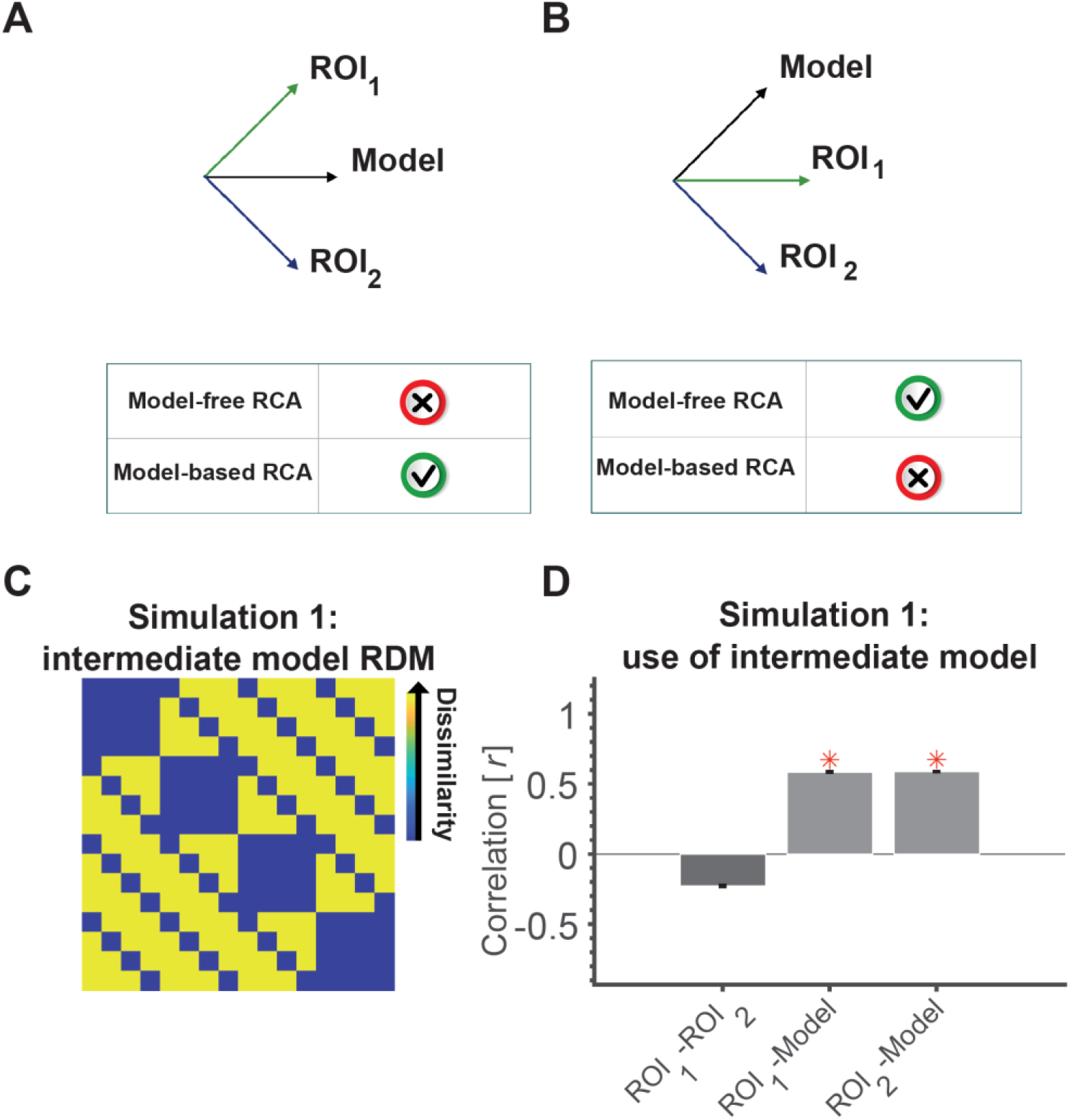
An intermediate model RDM can make two uncorrelated ROIs (RDMs) look connected. RDMs can be represented as a unit vector emanating from the origin in the N-dimensional space (N = number of elements in the RDM, i.e. n(n-1)/2 with n being the number of conditions). (A) Shows a situation where a model RDM has equal angle to two orthogonal (unrelated) neural RDMs. Looking from the lens of this model can leave the impression that the two neural RDMs are similar as they project similarly on the model RDM. (B) Shows a situation where two neural RDMs are positively correlated, but might look unrelated (disconnected) when looking at them through the lens of the model RDM as only one has a component along the model RDM’s vector and the other is orthogonal. (C) The model RDM used in Simulation 1, which has components of the representations in ROI 1 (position) and ROI 2 (semantic category). This model RDM has equal correlations (= 0.6) to the models used to generate neural RDMs of the two ROIs (shown in Figure 1, top); almost similar correlations to the neural RDMs (0.6). (D) The correlation between the neural RDM of the two ROIs (model-free RCA) and between the neural and model RDMs of each ROI (model-based RCA). Red asterisks show significant above-chance correlation values (connectivity) as evaluated by a one-sided Wilcoxon’s signed rank test against zero.

These examples show that model-based and model-free RCA can lead to inconsistent results. One might think that this is a consequence of using Pearson correlation or correlation-based measures for comparing RDMs. However, it is easy to imagine cases where using other metrics for comparing RDMs also leads to similar dissociations. The fact that model-free RCA is based on direct comparison of neural RDMs in the original dissimilarity space and that model-based RCA is based on comparison of projected RDMs on a line (projection) defined by the model, means it is always possible that the two will give inconsistent results. The degree of inconsistency would depend on the neural RDMs and the direction defined by the model. It might be worth noting that while there have been other implementations of model-based RCA (e.g. Karimi-Rouzbahani et al., 2021a; Clarke et al., 2018), they all assess similarity of RDMs along projections. Therefore, caution is needed.

Next, we consider three different scenarios, in which model-based and model-free RCA give inconsistent results. First we show how the incorporation of model(s) in model-based RCA can provide different results to model-free RCA. To that end, we present a scenario (similar to Figure 2A) where applying model-based RCA to two ROIs that represent independent aspects of visual stimuli results in the wrong conclusion that the two ROIs have shared information.

The general settings are explained in section *Simulations*. We used a model RDM that incorporates both aspects of the stimuli (i.e. position and object category). This intermediate model hypothesized a larger pattern dissimilarity for two conditions that are different in both position and object category. This intermediate model showed equal level of correlation/similarity to the neural RDM in each of the two ROIs (r = 0.6 between the model in Figure 2C and the RDMs shown in top and/or bottom panels of Figure 1). To implement model-based RCA, we simply calculated the correlation between the neural RDMs and the model and statistically compared the results across ROIs. To implement the model-free RCA, we calculated the correlation between the neural RDMs of the two ROIs directly (Basti et al., 2020; Karimi-Rouzbahani et al., 2021c; Shahbazi et al., 2021).

Clearly two ROIs which encode/represent statistically unrelated information are not connected. However, looking at the two ROIs through the lens of an intermediate model in model-based RCA can leave the impression that the ROIs are connected. Model-based RCA (i.e. the correlation between the model and each of the ROIs) showed significant positive correlation for two ROIs with the same model. The correlations between the model and the two ROIs were not statistically different (Wilcoxon’s signed rank test; p = 0.94). This (incorrectly) suggests that the two ROIs are connected (Figure 2D). However, model-free RCA (i.e. direct correlation between the two ROIs) correctly showed no positive correlation between the ROIs suggesting no connectivity. It might be worth adding that had we used a simple model instead of the intermediate model, for example one of the two models illustrated in the top panel of Figure 1B, we would have correctly observed no model-based RCA. Therefore, the issue relates to the representational structures of the two ROIs as well as the model used for examining their connectivity.

One way to avoid the false connectivity observed here for model-based RCA is to try to use minimal models where only one, rather than several, aspects of information is tracked between ROIs. The fact that our intermediate model had components from both aspects of stimuli (i.e. position and category) made it possible to capture variances explained by different processes, i.e. independent encoding of each aspect. Using simple models, for example models that correspond to simple hypotheses, enables untangling representational connectivity along different dimensions of information transfer. However, it might be difficult to come up with simple models in most cases, unless the underlying representations are already well characterised.

It is of note that, while we implemented a simplified version of model-based RCA in Simulation 1, some implementations in the literature have incorporated other parameters, such as time and delay, and other techniques such as multi-linear regression and partial correlation (Goddard et al., 2016; Karimi-Rouzbahani et al., 2018; Karimi-Rouzbahani et al., 2019; Goddard et al., 2019; Karimi-Rouzbahani et al., 2021a) each of which may affect the results. However, as the problem addressed here is a general issue, it may also be a concern for other implementations.

### 2.2 Simulation 2: spurious connectivity from common patterns across regions with distinct representations can be avoided using model-based RCA with proper models

There can be situations where common uninformative patterns are present along with the informative representations in the pair of ROIs considered for connectivity. The common patterns can be as simple as measurement or neural noise which might be statistically dependent across areas and/or the leakage or feeding of uninformative activations from a third ROI to both ROIs as a result of proximity and/or poor spatial resolution (e.g., in EEG and MEG). On the other hand, it can also be the case that the two ROIs encode/represent some shared aspects, which are either task-irrelevant or not the target of study. These aspects might be independently represented in the two ROIs under study, or be inherited from a third ROI. For example, both position-selective early visual area (V2) and the semantically selective area (ITC) can be sensitive to low-level image statistics such as the spatial frequencies of the stimulus due to connections from V1, which can lead to false/spurious connectivity if their RDMs are directly compared (as in model-free RCA). This might be because both the V2 and ITC RDMs also contain spatial frequency information. Note that the reason we consider representational connectivity in these cases to be false positive is that we are not interested in capturing commonality in noise, or in capturing this low level information (i.e. spatial frequency) which are represented in both ROIs, but rather by the particular information for which we have hypotheses.

Below we simulate the impact of adding common patterns of activation to a pair of ROIs which otherwise represent distinct information, and show how model-free RCA, and some implementations of model-based RCA, can be affected. We show that using proper models that match the dominant representations of the two ROIs can mitigate the false connectivity.

The neural patterns generated here are the same as Simulation 1 (with no connectivity between the two ROIs) except that now we also include the time course of representations to be able to implement more realistic model-free and model-based RCAs (rather than the simplified ones implemented in Simulation 1). We added the temporal dimension so that correlations could be computed over time. ROI representations at different time points were consistent with the same structures depicted in Figure 1B. In other words, the information was represented constantly over time but experienced some additive Gaussian noise. We simulated the activity patterns of the two ROIs over 200 time samples. The two ROIs were assumed to encode the two above-mentioned distinct aspects of information (i.e. position and semantic categories).

We performed model-free RCA by calculating the direct correlation between RDMs of the two ROIs at every time point and then averaging the resultant correlations over the simulated time window.

In this Simulation (and also the next Simulation), we consider two versions of model-based RCA. Generally, to perform model-based RCA, we first obtain the correlation between the neural and the corresponding model RDM of each ROI on every time point and then calculate the correlation between the time courses of neural-model correlations for the two ROIs. In one version, we consider one common model for the two ROIs (similar to Simulation 1) and in the other we use ROI-specific models (i.e. one model per ROI). The reason behind using the 1-model RCA is that the experimenter might simply want to evaluate potential information exchange between areas about a specific known aspect of information (e.g. familiarity information across occipital vs. frontal areas: Karimi-Rouzbahani et al., 2021a). On the other hand, the experimenter might have hypotheses about the dominant aspect of information represented in each of the two ROIs which might be different (e.g. visual information in lower visual areas vs. semantic information in ITC: Clarke et al., 2018). In this case it might be suitable to compare each ROI to a specific model of itself and use a 2-model RCA. The interpretation of 2-model RCA would be different and will be explained in Simulation 3. Accordingly, when using the 1-model RCA, we used the position model for both ROIs despite the fact that ITC represented category information. For the 2-model RCA, we used the position model for detecting position information in ROI 1 and a semantic-category model for detecting semantic-category information in ROI 2; therefore, the models perfectly matched what was dominantly represented in each ROI (see Figure 1).

We added a non-structured (noise) pattern to both ROIs and evaluated its impact on connectivity. To generate the common pattern, we used Gaussian noise and a random transformation matrix to impose correlated noise across areas. Here, we can assume that the two ROIs that encode/represent independent aspects of the stimuli are not connected. However, we show that the common pattern can lead to false/spurious connectivity in model-free and 1-model RCA.

Before adding the common patterns to the ROIs, the three connectivity measures were either negative (model-free RCA) or around-zero (1-model and 2-models RCA), suggesting no connectivity between the ROIs (Figure 3A). However, the addition of common patterns to the two ROIs led to spurious connectivity for model-free and 1-model RCA, with both showing significantly above-chance RCA. This was predictable for the model-free RCA because it relies on the representations of information in ROIs, which become correlated by the added correlated noise.

**Figure 3.**
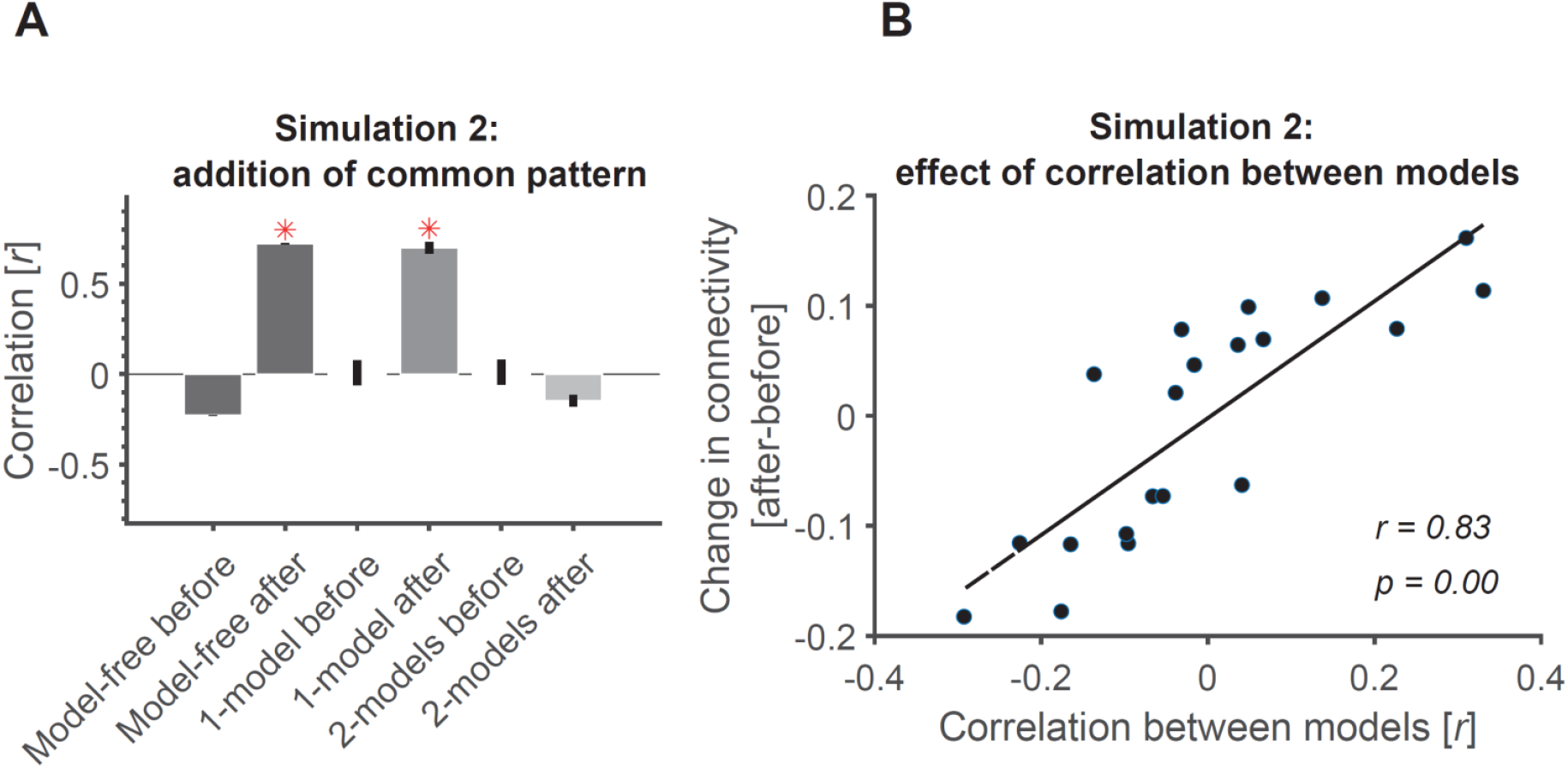
Addition of common pattern to pair of ROIs can make them look connected in model-free and 1-model RCA but not 2-model RCA if the ROIs are distinct enough. (A) Addition of common non-structured (noise) patterns to both ROIs leads to significant connectivity when evaluated using model-free and 1-model RCA, but not 2-model RCA because the two ROIs dominantly represent information which are negatively correlated (See Figure 2). Red asterisks show significant above-chance correlation values (connectivity) as evaluated by one-sided Wilcoxon’s signed rank test against zero. (B) The more distinct the dominant information represented across the ROIs, the less the effect of added common noise on their connectivity. Dots show the amount of change in connectivity as a function of correlation between the information represented in the two model RDMs used in 2-model RCA. Each dot represents data from a single simulated subject. The line shows the best linear fit to the data. The correlation and the significance of correlations are also shown as calculated using Pearson’s linear correlation.

This result was especially interesting for 1-model RCA because the method required the two ROIs to be temporally correlated to show connectivity. This confirms that the common pattern has not only correlated the representations of the two ROIs, but it has also added temporal correlations to the representations of the two ROIs making them fluctuate similarly over time (which is key for our model-based measure). We also observed that this spurious connectivity for 1-model RCA remained when using any arbitrarily defined random models. Specifically, we observed that even models unrelated to those relevant to the ROIs (e.g. position and/or semantic categories) could lead to false connectivity in 1-model RCA. This can be explained by the fact that the two correlated representations in the ROIs, as imposed by the added noise pattern, will lead to similar patterns of correlation to any common model.

Finally, despite the correlations imposed on the contents of representations and the temporal patterns across ROIs, the 2-model RCA (correctly) showed negative correlations between the ROIs suggesting no connectivity. This negative correlation/connectivity can be explained by the fact that the two correlated representations will negatively correlate when evaluated against two negatively correlated RDMs. This result suggests that 2-model RCA is influenced by the relationship between two ROIs as reflected in their optimal model RDMs. More specifically, it seems that if two ROIs’ model RDMs correlate negatively (suggesting distinct codes represented in each of them), they can remain immune to the added common noise. To test this hypothesis, we generated 2 random models the two ROIs of each simulated subject and calculated the level of decline in correlation/connectivity from before to after adding the common noise. Results showed that the less similar the models (and the corresponding ROIs), the less the value of connectivity under the influence of the added noise (Figure 3B). In other words, correlated added patterns will be more likely to lead to spurious connectivity if the two ROIs represent similar information and have a larger overlap.

Therefore, in our case where our two ROIs originally represented two distinct information, we remained unaffected by the added common noise simply because the chosen model RDMs which represented the dominant aspect of information in each ROI were not positively correlated. However, the choice of a single model in such situation can lead to false connectivity, as was the case for 1-model RCA.

### 2.3 Simulation 3: model-based RCA detects transformation of information by testing the relationship of temporal dynamics of different types of information across ROIs

There can be situations where the structure of the information is transformed from one ROI to the next. In fact, it seems unlikely that information remains untransformed (“copied”) between any two ROIs in the brain. Therefore, direct comparison of neural representations, as implemented in model-free RCA, can miss such potential connectivity simply because the statistical relationship may be lost in transformation. However, model-based RCA may allow us to detect the connectivity between two areas which encode distinct information based on their temporal statistical congruency. Below we simulate two ROIs that represent two distinct aspects of information, with dynamics that are either temporally congruent/incongruent between ROIs. Specifically, the information about the stimulus position initially appears in the source ROI (V2) and is followed by the semantic-category information which appears in the destination ROI (ITC)^2^. This scenario resembles a study which showed causal information-transfer and transformation from early visual to ITC areas (Clarke et al., 2018). As the detection of connectivity using model-based RCA with ROI-specific models also needs the adoption of correct model for each ROI, because the information is transformed, we also check the effect of choosing the correct model for each ROI.

We used simulations to investigate this feature of model-based RCA. The details of information representation in the two ROIs in this simulation are identical to Simulation 2, with the exception that the information does not appear throughout the simulation window but rather for a fixed period of time in each ROI (samples 30 to 60 in ROI 1; Figure 4A). There was a delay of ±20 samples between ROIs 1 and 2 (positive for congruent and negative for incongruent case) which was jittered between 0 to 10 samples (uniform random distribution) across the simulated participants (N = 20). This led to information appearing in ROI 1 before ROI 2 in congruent cases, and in ROI 2 before ROI 1 in incongruent cases (Figure 4A). Specifically, patterns could appear between samples of 40 to 80 in ROI 2 in the congruent case and between samples of 0 and 40 in ROI 2 in the incongruent case. The activity patterns of the two ROIs did not contain any information in the samples outside the mentioned windows. Note that the information which was dominantly represented in the two ROIs differed as in the above simulations (position encoding in V2 (source) and semantic category encoding in ITC (destination)). This scenario simulated information flow from the source to the destination area that has been evaluated in previous studies using both model-free and model-based RCA (Goddard et al., 2016; Karimi-Rouzbahani, 2018; Clarke et al., 2018; Karimi-Rouzbahani et al., 2019; Goddard et al., 2019; Karimi-Rouzbahani et al., 2021a). In this analysis, the onsets of information appearance predict the direction of information transfer (e.g. potential information flow from V2 to ITC). Here we only report the connectivity from ROI 1 to ROI 2 (feed-forward information flow in the ventral visual stream) and not vice versa. In testing the connectivity for both model-free and model-based RCA methods, we set the analysis delay time (i.e. lag) between ROIs to be 20 (no jitter) for all our participants. This parameter is usually set by the researcher and fixed across participants (Goddard et al., 2016; Karimi-Rouzbahani, 2018; Karimi-Rouzbahani et al., 2019; Goddard et al., 2019; Karimi-Rouzbahani et al., 2021a). We performed model-free RCA by calculating the direct correlation between the RDM of the source ROI at time *t* and the RDM of the destination ROI at time *t* + τ where τ refers to the delay (= 20 samples) and then averaged the time course of correlations within each subject. For model-based RCA, we calculated the correlation between the neural and model RDMs for each ROI on every time point as in Simulation 2 (note that we considered the two cases of having one model RDM or two different model RDMs), shifted the model-correlation time course of ROI 2 by 20 (jittered between 0 to 10w samples) relative to ROI 1, and computed their correlation coefficient. In both model-free and model-based analyses, the incorporation of the delay led to compensating for the inter-ROI delay in the data.

**Figure 4.**
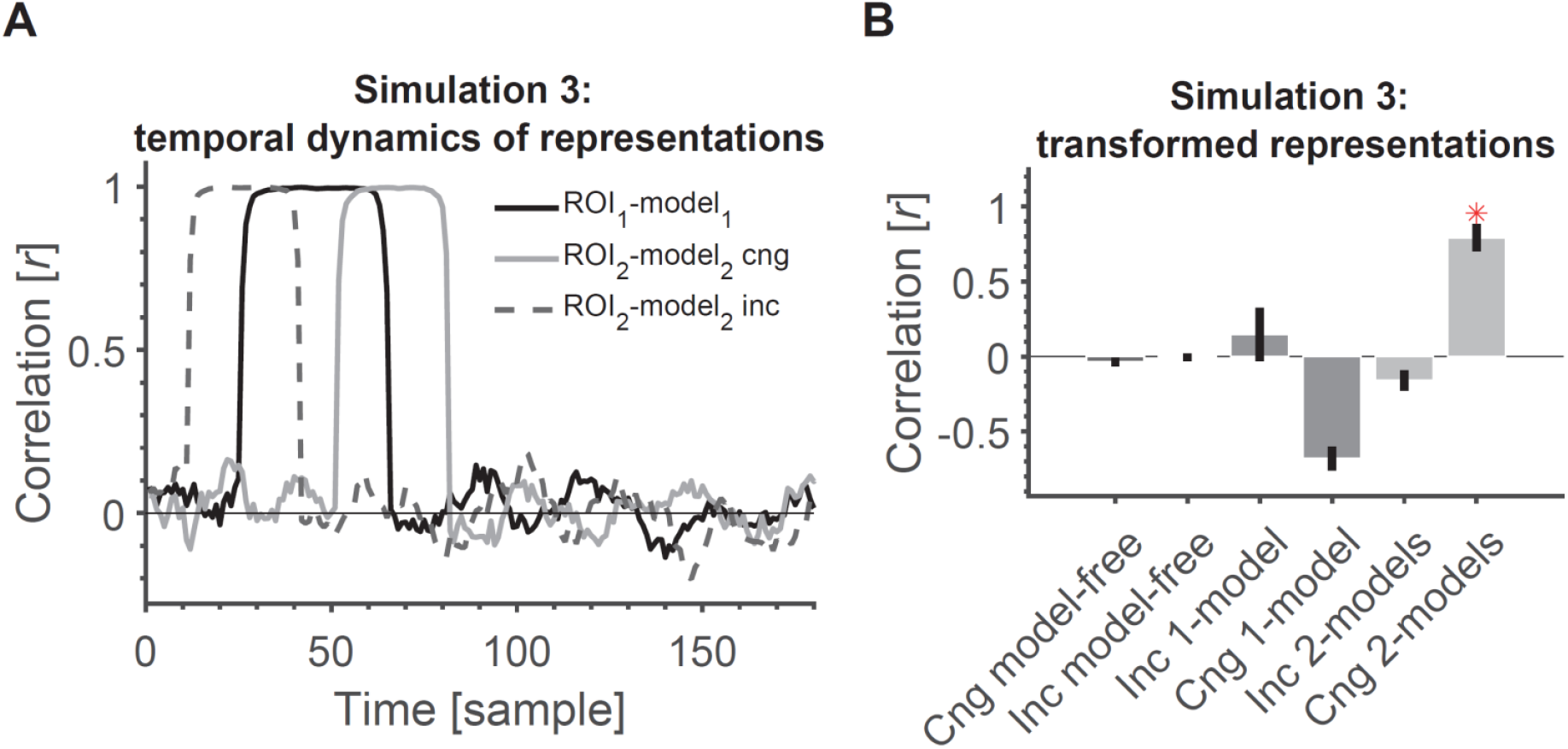
2-model model-based RCA allows us to detect transformation of information if the temporal dynamics across ROIs are statistically related/congruent. (A) Time course of information encoding in the two ROIs with at a delay of 20 samples (congruent (cng), solid grey line) and a delay of −20 from ROI 1 to ROI 2 (incongruent (inc), dashed grey line). The delay was variable across participants. The time courses show the correlation between each ROI and its corresponding model (position model for ROI 1 and semantic-category model for ROI 2). (B) Transformation of information across ROIs causes all model-free and model-based RCAs to miss the connectivity except when the information appears congruently across ROIs (first in ROI 1 followed by ROI 2) and using 2-model RCA. Red asterisk shows significant above-chance correlation value (connectivity) as evaluated by one-sided Wilcoxon’s signed rank test against zero.

When applying this to real data, it is important to have a reasonable estimate of the delay. This could correspond to the physiological delay. A more principled way would be to partition the data and estimate the optimal delay from one half and apply it to the other half. However, this requires independent measurements of the same task in each subject. A more extended version of the RCA could be to perform Granger causality to examine Granger-causal relationships between areas as in previous studies (Goddard et al., 2016; Karimi-Rouzbahani, 2018; Clarke et al., 2018; Karimi-Rouzbahani et al., 2019; Kietzmann et al., 2019; Goddard et al., 2019). That would also be subject to similar considerations. However, comparing the different approaches at a conceptual and mathematical level is beyond the scope of the current study.

Our assumption here is that the two ROIs that encode/represent statistically unrelated information can be considered connected if their temporal information encoding profiles are statistically related/congruent (representations appear in the destination after the source ROI at around the hypothesized delay). However, such relationship might not be detected using model-free RCA or model-based RCA with a common model.

Simulation results show that model-free RCA did not detect any connectivity between the two ROIs (Figure 4B). The 1-model RCA also failed to detect the connectivity whether the representations in ROIs appeared congruently and incongruently. The reason is that the representations that were transformed from the source to the destination ROI no longer matced the common model in the destination ROI (we used the model RDM from ROI 1 for both ROIs). The 2-model RCA also failed to detect the connectivity when the representations appeared incongruently across the ROIs (first in the destination followed by the source) because information time courses in one did not reliably follow the other according to the hypothesized lag. Finally, the 2-model RCA could detect the connectivity when the representations appeared congruently across the ROIs. Therefore, for the transformed information to be detected, one needs to have both accurate models of information representations as well as correct prior knowledge about temporal dynamics of information across ROIs.

Note that in these simulations, we incorporated the delay in our analysis and the two ROIs followed the temporal profiles of representations shown in Figure 4A. Therefore, the absence of connectivity in the model-free and 1-model RCA cannot be explained by the fact that we used lagged correlations in the 2 model case. To this end, we incorporated the delay in all our RCA measures here.

Model-free RCA is only sensitive to direct statistical relationship between neural RDMs across time, and fails to detect the connectivity if the two ROIs do not statistically relate. However, the 2-model RCA implemented here allows for the detection of congruent inter-ROI statistical dependencies by having models that capture the representational structure of each ROI. Interestingly, 2-model model-based RCA can also detect the transformation of information that is not inherited from confounders. Similar to the observation made in Simulation 2, it might be that common task-irrelevant patterns in both ROIs obscure the shared information as captured by 1-model RCA or the transformation of information as captured by 2-model RCA. A solution to this, which comes at the expense of knowing the structure of the common patterns, would be to remove their contribution by regressing out the RDM of the common pattern from the RDM of each ROI at each time-point.

One solution to the above failure of model-free RCA to detect connectivity might be to use non-linear mapping functions. Such functions allow more flexible relationships to be detected between areas despite drastic transformation of the representational structure. Such non-linear mapping functions include distance correlation (Geerligs and Henson, 2016), projection to a Riemannian manifold (Shahbazi et al., 2021) or more general functions estimated by artificial neural networks (Anzellotti et al., 2017b). These potential solutions are not investigated here.

## 3. Discussion and Conclusions

Multi-dimensional connectivity is a rapidly developing area of brain connectivity analysis. One of the approaches to multi-dimensional connectivity is representational connectivity analysis (RCA). RCA quantifies the similarity of inter-relationship between the neural representations across experimental conditions for distributed patterns of activity of two brain regions (Kriegeskorte et al., 2008). This allows us to track “information” (by the representational geometry in the multi-dimensional response space) rather than the mere similarity of average response levels across two regions. Despite its versatility, a better understanding of the situations that can challenge and/or mislead RCA is needed. In this manuscript, we explain two main approaches of RCA. One is model-free RCA that directly compares the representational geometries of two brain ROIs. Model-free RCA can tell us whether two brain ROIs have any shared information in their multi-dimensional response patterns. The other is model-based RCA. In this paper, we make a distinction between two approaches to model-based RCA. We think this distinction is important, since besides the difference in technical details and implementation, they entail different interpretations about regional interactions. One variant of model-based RCA, which uses a common model, tests whether the representational geometries of the two ROIs are similarly concordant to a hypothesized geometry (i.e., the model). This can tell us whether two brain regions have shared information with regard to a specific aspect of the stimuli/task. The other variant of model-based RCA uses ROI-specific models, which, with time-resolved data, tells us whether information in one region is transformed into different information in another region. Therefore, while this also pertains to functional connectivity, it does not explicitly get at shared information.

Model-free and model-based RCA can potentially provide inconsistent results in certain circumstances. These inconsistencies depend on many factors, some of which are the spatiotemporal structure of neural representations and the choice of the model(s) used in the analysis. Here, we focused on three scenarios where model-free and model-based RCA provided opposing connectivity results.

In the first scenario, we simulated a situation where the neural representations across a pair of regions showed unrelated information. As expected, model-free RCA showed no connectivity between the pair of regions. Interestingly, however, we observed that using model-based RCA with an *intermediate* model, which contains information about the representations in both regions, can leave the false impression that the two regions are connected. Specifically, the two regions showed almost equal, positive and significant correlation to the intermediate model suggesting that from the “lens” of the selected model, the two regions appear to be connected.

There are three considerations about this simulation. First, although for simplicity we did not directly implement either of the two published methods of model-based RCA (Clarke et al., 2018; Karimi-Rouzbahani et al., 2021a), the problem we pointed out here can affect both those methods. This is because they compare the correlation between the models either explicitly (Clarke et al., 2018), or implicitly within the formulation of partial correlation (Karimi-Rouzbahani et al., 2021a). Second, although we used a two-component model for this simulation to simplify the interpretation, this situation is not limited to two-component models. In fact, any other models that would share positive and relatively equal amount of variance with different components encoded in two areas could be used for the first scenario and would lead to similar interpretations. Third, the false connectivity observed in this scenario is not driven by the specific similarity metric we used (i.e. Pearson’s linear correlation). Although different similarity metrics show different characteristics (Walther et al., 2016; Shahbazi et al, 2021), as long as the selected metric provides similar values for the similarity between two different neural and a given RDM model, a similar effect will be observed. The reason is that all similarity metrics summarize a high-dimensional representational space into a single-dimensional space, which inevitably leads to loss of information. Finally, on the other end of the spectrum, there can be cases where two regions represent one or several very similar aspects of information, but they still look unrelated/disconnected through the lens of a particular model. However, this case seems less problematic since the main reason behind using model-based rather than model-free RCA is to limit the representations to the desired information (Karimi-Rouzbahani et al., 2021a). In fact, with model-based RCA, similarity is intended to be evaluated through the lens of model(s) and that might be desirable in many scenarios. Therefore, it would be good practice to do perform both types of RCA together with RSA (information mapping) while being aware of the limitations and caveats of each.

In the second scenario, we simulated a situation where the addition of statistically-related patterns of activity to a pair of statistically-unrelated regions imposed a statistical relationship between. This lead to apparent connectivity in model-free RCA and when using 1-model RCA. However, the common pattern did not affect apparent connectivity when using 2-model RCA, as long as the two models were orthogonal and the two ROIs represented distinct information. Please note that the added common pattern can be non-structured or structured. Although we have observed that both common noise (non-structured) and structured patterns (data not shown) led to similar results, the structure of the common pattern can affect the connectivity as a result of interaction with the representations in the regions and models. It is also of note that the addition of common patterns does not always inflate the connectivity (e.g. in model-free or 1-model RCA); it can also decrease it leading to missing the connectivity. For example, if two regions are perfectly correlated, the addition of common noise (if not perfectly identical but only statistically related across regions) could lead to a decline in model-free RCA as a result of distorting the patterns. Generally, both model-free and model-based RCA can be affected by the noise as a result of the complex interaction between the representation in each region, the structure of the added pattern, the models, and the temporal dynamics of representations. Therefore, despite the situation shown in Simulation 2, these methods we are still far from remaining immune from common noise. We can, however, understand where we are most susceptible to it. One simple remedy for the effect of common patterns (as long as the common pattern is additive), would be to regress out its contribution from the RDMs of the two ROIs prior to computing connectivity measures. This is in spirit similar to our recent implementation of model-based RCA (using partial correlation), where we partialled out the effect of additional low-level image statistics from the two regions under study (Karimi-Rouzbahani et al., 2021a).

In the third scenario we simulated a situation where two regions encoded different types of information that were either temporally incongruent or congruent. In other words, the information initially appeared in one region and after some delay in the other region (temporally congruent). Model-based RCA with proper choices of models can capture the relationship between the different representations. The transformation of information seems an integral part of brain connectivity as it seems unlikely that information would remain intact from one brain region to another (Lahaye et al., 2003; Hlinka et al., 2011). Transformations of information have already been reported in visual system of human and monkey brain (Dicarlo et al. 2012; Kietzmann et al., 2019) and are implemented by other sensory hierarchies as well (Winkowski and Kanold, 2013). For example, it has been suggested that visual information is moved from low-to a high-dimensional space along the ventral visual stream and brought back to the low-dimensional space in later stages of the stream to compensate for variations of visual objects and form semantically-categorized object clusters (Dicarlo et al., 2012; Karimi-Rouzbahani et al., 2017a; Karimi-Rouzbahani et al., 2017b). Using model-based RCA, previous work has found that information transforms from visual to semantic brain areas (Clarke et al., 2018). The delay in the analysis also potentially captures the neural lag in information transfer in the brain (Cichy et al., 2014). Despite the fact that the delay was incorporated in all the different connectivity analyses, the drastic transformation of information simulated in Simulation 3 led to the connectivity being missed by model-free RCA. In addition, this simulation also pointed out the importance of the delay in connectivity analysis matching the data. The delay is generally set *a priori*, meaning that choice of improper delays (negative vs. positive; which also determines the direction of information) can lead to missing the connectivity. Despite these details, 2-model model-based RCA detected the connectivity as a result of its simultaneous sensitivity to targeted region-specific information representation and the temporally congruent patterns of information representation. Therefore, a hypothesis-driven method of RCA allows us to detect information that is transformed as it passes between brain regions.

It is generally desired that a connectivity method determines the transferred content, direction and temporal dynamics of information flow when possible (e.g. in M/EEG). To that end, previous studies implemented techniques including partial correlation (Goddard et al., 2016; Karimi-Rouzbahani, 2018; Karimi-Rouzbahani et al., 2019; Goddard et al., 2019; Karimi-Rouzbahani et al., 2021a) and regression (Kietzmann et al., 2019), or tested for Granger causal relationship between areas (Goddard et al., 2016; Clarke et al., 2018; Karimi-Rouzbahani, 2018; Karimi-Rouzbahani et al., 2019; Goddard et al., 2019), or used models to measure the contribution of one area to another in the direction of the task (Karimi-Rouzbahani et al., 2021a) or incorporated autoregressive approaches to estimate proper delay between areas (Clarke et al., 2018). In our most recent effort, to bring together the advantages of the mentioned methods, we proposed a variant of model-based RCA which provided information about the *content* of the transferred information, its *direction* and *temporal dynamics* simultaneously (Karimi-Rouzbahani et al., 2021a). This method showed distinct dynamics and direction of face familiarity-information flow across peri-frontal and peri-occipital cortices for different levels of perceptual uncertainty. Despite our minimalist approach in the current study, the insights and cautions provided by this work can be generalized to more complex implementations of RCA as well.

Additionally, one could also consider other extensions to model-free RCA. Similar to “information connectivity” (Coutanche, 2013) where multi-dimensional connectivity is established by correlating time series of classification-accuracies across regions, one can compare time courses of the exemplar discriminability index (EDI, Nili et al., 2020) across regions. EDI is a model-free RSA statistic in each region and quantifies the extent to which different experimental conditions elicit distinct patterns of activation. Similar to the implementation of model-free RCA, this definition of model-free RCA also does not shed light into the content of shared information.

A limitation of the current study is that we only evaluated connectivity using linear, rather than non-linear, relationships. While this simplification allowed us to make more intuitive predictions about the relationship between brain responses and the models, a more general approach would be to incorporate non-linear connectivity between areas as well. While we believe that the cases evaluated in Simulations 1 and 2 will not be affected by using a non-linear connectivity metric, non-linear mapping functions in Simulation 3 (Geerligs and Henson, 2016; Anzellotti et al., 2017b; Basti et al, 2020; Shahbazi et al; 2021) may allow for detecting non-linear relationships between areas. Therefore, future studies will need to evaluate the impact of non-linear mapping functions in RCA.

This work takes initial steps towards better characterization of the model-free and model-based RCA approaches that have been increasingly used in recent years. We tried to make the scenarios as general as possible, so that the insights can be generalized to different implementations of the two general classes of model-free and model-based RCA. Therefore, the points made here can provide insight when studying brain connectivity using variety of neural recording modalities such as EEG, MEG, multi-electrode electrophysiology and fMRI.

## Acknowledgments

This research was funded by UK Royal Society’s Newton International Fellowship NIF\R1\192608 to H.K.R. and MRC intramural funding SUAG/052/G101400 and SUAG/046/G101400 to A.W. and R.H., respectively.

## Footnotes

1 We use the term “model” in a general sense: it can be a conceptual model, a computational model or a third brain region, etc.

2 We consider the case where the information is reliably transferred from one ROI to another as temporally congruent and cases where there is no transfer of information or the transformation is not reliable/consistent as temporally incongruent.

